# Differentiating BOLD and non-BOLD signals in fMRI time series using cross-cortical depth delay patterns

**DOI:** 10.1101/2024.12.26.628516

**Authors:** Jingyuan E. Chen, Anna I. Blazejewska, Jiawen Fan, Nina E. Fultz, Bruce R. Rosen, Laura D. Lewis, Jonathan R. Polimeni

## Abstract

Over the past two decades, rapid advancements in magnetic resonance technology have significantly enhanced the imaging resolution of functional Magnetic Resonance Imaging (fMRI), far surpassing its initial capabilities. Beyond mapping brain functional architecture at unprecedented scales, high-spatial-resolution acquisitions have also inspired and enabled several novel analytical strategies that can potentially improve the sensitivity and neuronal specificity of fMRI. With small voxels, one can sample from different levels of the vascular hierarchy within the cerebral cortex and resolve the temporal progression of hemodynamic changes from parenchymal to pial vessels. We propose that this characteristic pattern of temporal progression across cortical depths can aid in distinguishing neurogenic blood-oxygenation-level-dependent (BOLD) signals from typical nuisance factors arising from non-BOLD origins, such as head motion and pulsatility. In this study, we examine the feasibility of applying cross-cortical depth temporal lag (CortiLag) patterns to automatically categorize BOLD and non-BOLD signal components in modern-resolution BOLD-fMRI data. We construct an independent component analysis (ICA)-based framework for fMRI de-noising, analogous to previously proposed multi-echo (ME) ICA, except that here we explore the across-depth instead of across-echo dependence to distinguish BOLD and non-BOLD components. The efficacy of this framework is demonstrated using visual task data at three graded spatiotemporal resolutions (voxel sizes = 1.1, 1.5, and 2.0 mm isotropic at temporal intervals = 1700, 1120, and 928 ms). The proposed CortiLag-ICA framework leverages prior knowledge of the spatiotemporal properties of BOLD-fMRI and serves as an alternative to ME-ICA for cleaning moderate- and high-spatial-resolution fMRI data when multi-echo acquisitions are not available.

## 1 Introduction

High spatial resolution is a key feature that makes functional Magnetic Resonance Imaging (fMRI) one of the most widely-used non-invasive techniques for experimental human neuroscience. The rapid advancement of MR engineering over the past two decades has pushed the imaging resolution of fMRI to be far higher than what it was at inception. For instance, using Simultaneous Multi-Slice imaging (a.k.a. “multiband” echo-planar imaging or EPI) (Barth et al., 2016; Feinberg et al., 2010; Feinberg and Setsompop, 2013; Feinberg and Yacoub, 2012; Hennig et al., 2007; Lin et al., 2006; Moeller et al., 2010; Narsude et al., 2016; Setsompop et al., 2016, 2012; Zahneisen et al., 2011), several major, large-scale neuroimaging studies have collected and shared 3T whole-brain fMRI datasets with ∼2-mm isotropic voxels sizes and sub-second repetition time (Bookheimer et al., 2019; Casey et al., 2018; Miller et al., 2016; Van Essen et al., 2013); and—when combined with ultra-high magnetic field scanners, advanced instrumentation, and parallel imaging strategies—state-of-the-art fMRI is able to achieve voxel sizes down to the sub-millimeter scale (Berman et al., 2021; Feinberg et al., 2023, 2018; Han et al., 2021; Huber et al., 2017; Vizioli et al., 2023). A compelling motivation for acquiring smaller voxel sizes is the potential for mapping the brain’s functional architecture at fine spatial scales, achieved by localizing hemodynamic responses within subcortical and brainstem nuclei as well as cerebral cortical columns and layers (De Martino et al., 2013; Faull et al., 2015; Koopmans et al., 2010; Nasr et al., 2016; Polimeni et al., 2010; Satpute et al., 2013; Yacoub et al., 2007), thus yielding unprecedented insights into the brain activity underlying human cognition.

Accompanying the enthusiasm for high-resolution fMRI acquisitions are new challenges in denoising these data, especially the removal of structured noise sources, as denoising approaches suitable for conventional resolutions may not extend well to high-resolution data (Polimeni et al., 2018; Wang et al., 2022). For instance, multi-echo independent component analysis (ME-ICA) (Evans et al., 2015; Kundu et al., 2017, 2013, 2012; Lynch et al., 2020; Olafsson et al., 2015), a robust and efficient method for addressing nuisance factors in conventional blood-oxygenation-level-dependent (BOLD) fMRI, is less accessible in high-resolution applications. This approach leverages the echo time (TE) dependence of BOLD contrast to distinguish functional information from noise, thereby enhancing the neuronal specificity of fMRI. Yet, incorporating multiple echoes in an fMRI acquisition typically incurs penalties in temporal resolution (due to the need to sample multiple echoes during the readout) or spatial resolution (as achieving increased image encoding typically requires reduced k-space coverage). Although high spatial resolution limits the feasibility of multi-echo acquisitions, it inspires new analytical strategies for denoising fMRI data. With small voxels, one can consider the cerebral cortex as a 3D volume instead of a 2D sheet, and sample from different levels of the cortical vascular hierarchy. Neurogenic BOLD changes are known to initiate within the parenchyma and drain outwards toward the pial surface (Tian et al., 2010), thus the ability to resolve this characteristic across-cortical temporal progression pattern offers a promising opportunity to distinguish functional signals from nuisance factors (such as head motion, cardiac pulsatility, etc.) in voxel-wise fMRI time series. See *Section 2 Theory* below for a detailed rationale.

In this study, we test the feasibility of differentiating between BOLD signal components and non-BOLD nuisance factors based on the temporal progression of BOLD-fMRI signals across cortical depths and establish an integrated, data-driven framework to automatically denoise moderate- and high-spatial-resolution BOLD-fMRI datasets. The framework is analogous to ME-ICA, except that here, we propose to examine the signal dependence across cortical depths instead of echoes. This article is organized as follows: we first introduce the rationale behind the idea of isolating functional BOLD signal components from non-BOLD artifacts according to the specific cross-cortical temporal lag patterns. We then introduce the overall scheme of the proposed framework and demonstrate its efficacy in enhancing the sensitivity of detecting visual task activation in gradient-echo (GE) BOLD-fMRI data at three graded spatiotemporal resolutions. Next, we show that integrating fMRI time series across voxels spanning the cortical thickness in an appropriate weighted average can produce activation maps with higher sensitivity than those based on data from any single cortical depth, analogous to optimized across-TE combination in ME-ICA. Finally, we discuss the pros and cons of the proposed de-noising framework relative to ME-ICA, and the feasibility of extending our denoising scheme to fMRI data acquired at lower spatial resolutions.

## 2 Theory

Because neuronal activity initiates the hemodynamic response within the parenchyma via neurovascular coupling (Hamel, 2006), and the BOLD signal propagates downstream along ascending venules toward the pial surface, BOLD time courses from superficial cortical depths lag in time behind those of the deeper cortical depths (Havlicek and Uludağ, 2020; Markuerkiaga et al., 2016). Empirical animal and human experiments have shown that the neurogenic hemodynamic lag between parenchymal and surface vessels is on the order of several hundred milliseconds (Siero et al., 2011; Tian et al., 2010) (illustrated in Fig. 1, see also Supplementary Fig. S1). Such cortical-depth-dependent temporal lag (CortiLag) patterns, by contrast, are not anticipated in major nuisance sources that confound fMRI signals. For instance, head movement-induced instantaneous intensity changes (from displacements of head position) and slower-scale intensity changes (from spin-history effects (Muresan et al., 2002)) should yield synchronous fMRI signal changes across cortical depths (except, perhaps, for special locations at which the slices run perfectly parallel to the cortex). Artifacts stemming from image acquisition or reconstruction such as Nyquist ghosting are also unlikely to exhibit specific spatial patterns that track the cortical geometry (Griffanti et al., 2017; Jezzard and Clare, 1999). As for quasi-periodical physiological artifacts time-locked to respiratory/cardiac cycles—including both magnetic field changes driven by changes in chest volume while breathing, and dynamic partial volume effects caused by pulsatile vessel/tissue displacement driven by the cardiac pressure wave—a zero or a minimal temporal lag, if it exists, would manifest across cortical depths. Therefore, CortiLag patterns provide an excellent opportunity to differentiate between neural activity and various forms of structured noise in BOLD fMRI time-series data.

**Figure 1:**
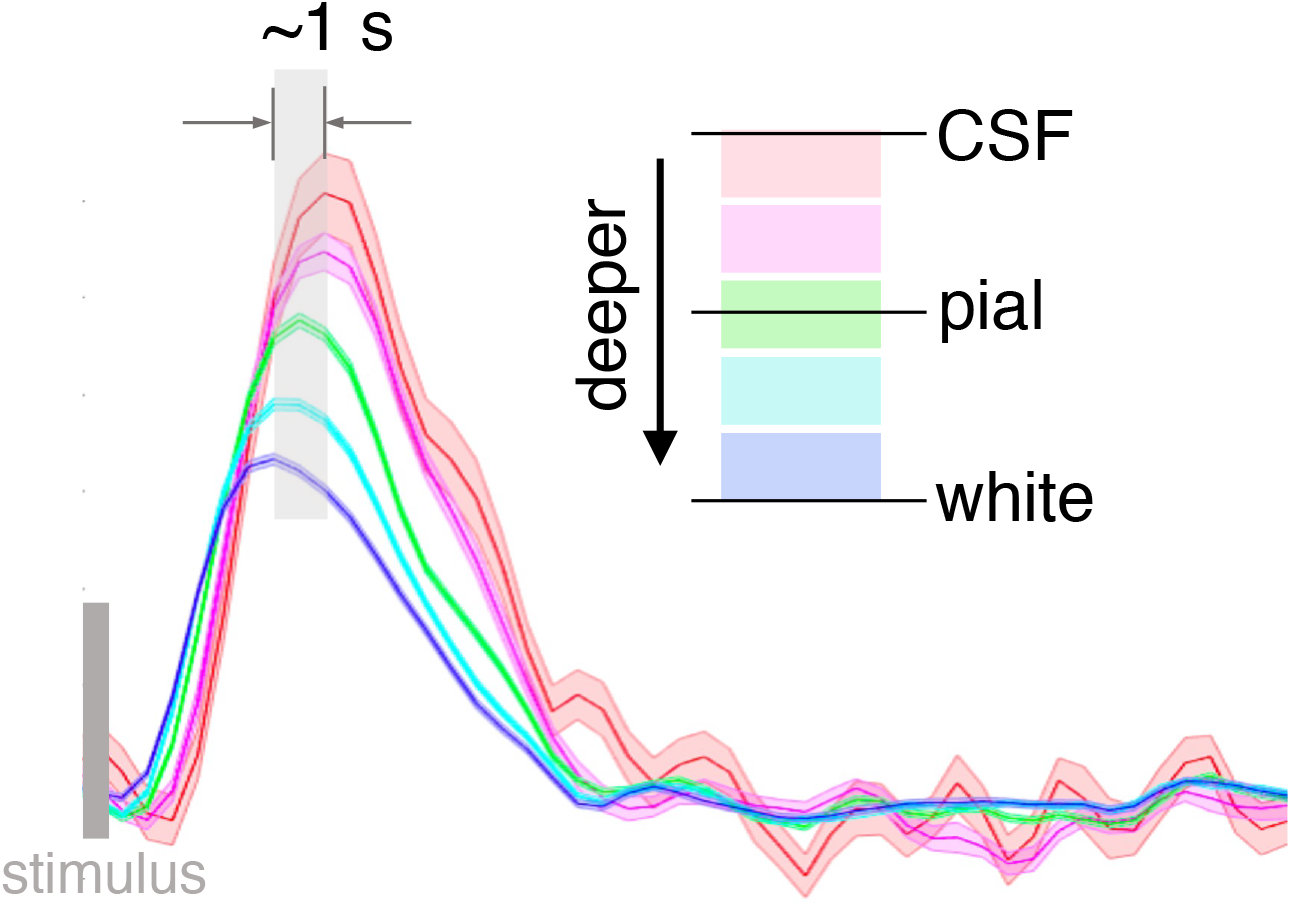
Neuronally-driven hemodynamic responses with graded temporal delays across cortical depths, illustrated using single-subject fMRI responses evoked by the flickering checkerboard visual stimulation (7T, 0.8-mm isotropic nominal voxel size, TR/TE = 1010/28 ms; data from study of Chen et al. (2021)). See Supplementary Fig. S1 for depth-dependent fMRI patterns generated using the 1.1-mm dataset from this study.

Given the limited spatiotemporal resolution of GE-BOLD fMRI, what are the lower bounds of temporal sampling intervals and voxel sizes needed to resolve differentiable CortiLag patterns between functional BOLD signals and non-BOLD noise? From a temporal-resolution perspective, as will be shown later in this article, even conventional sampling rates (e.g., ∼2 s) are sufficient to detect hemodynamic lags across cortical depths. As for spatial resolution, although it is unlikely to have multiple voxels spanning the entire cortical thickness at any given location with supra-millimeter resolutions, an approximately even distribution of voxel centroids across cortical depths can be achieved if a sufficiently large number of contiguous voxels are pooled (Polimeni et al., 2018). It is also of note that the across-depth distance between voxels exhibiting the earliest and latest BOLD responses can be larger than the cortical thickness (1–4.5 mm), due to the extravascular susceptibility field of large draining veins that can extend into subarachnoid CSF above the gray-pial border; hence voxel sizes as large as 2–3 mm isotropic, as employed in conventional fMRI studies, may suffice to dissociate earlier response in parenchyma from later responses observed in the CSF in most cortical areas. Taken together, although high-resolution acquisitions (e.g., approaching sub-second TRs or sub-millimeter voxel sizes) undoubtedly improve the detection of CortiLags, distinct CortiLag patterns between BOLD and non-BOLD signal components may also be detectable in moderate-resolution acquisitions (with 1–2 s TR and 2–3 mm iso. voxel size).

## 3 Methods

### 3.1 Volunteers and task

All experimental procedures were approved by the Massachusetts General Hospital Institutional Review Board. Twelve healthy subjects (24.7±4.3 years old, six females) were enrolled for this study, after providing written informed consent.

Each participant underwent three visual task scans with graded spatiotemporal resolutions (scan orders were counterbalanced across subjects). Visual stimuli were implemented using the MATLAB-based psychtoolbox (http://psychtoolbox.org), projected onto a screen mounted at the end of scanner bore, and viewed through a mirror fixed in front of the subjects’ eyes. A block-design paradigm was employed (task ‘on’: radial “checkerboards” counterphase flickering at 12 Hz; task ‘off’: gray background; 16/19 s on/off per block, 8 blocks per scan). To aid visual fixation and maintain vigilance, subjects performed a simple fixation task throughout the experiment. A small red dot at the center of the screen changed luminance randomly, with a mean interval of 2 s. Subjects were instructed to press a button upon detecting a luminance change, and their behavioral performance was continuously monitored. For all functional scans reported in this study, subjects achieved over 80% accuracy in detecting luminance changes in the fixation dot.

### 3.2 Acquisition

MR images were collected on a 7T Siemens MAGNETOM whole-body scanner (Siemens Healthineers, Erlangen, Germany) equipped with SC72 body gradients and an inhouse-built 32-channel brain receive-coil array (Keil et al., 2010). *High-resolution anatomical data:* High-resolution T_1_-weighted MPRAGE images (0.75-mm isotropic spatial resolution, with a 13-ms FOCI adiabatic inversion pulse (Zaretskaya et al., 2018)), TR = 2530 ms, TE = 1.76, 3.70 ms, flip angle = 7°, FOV = 240×240×168 mm^3^, bandwidth = 615 Hz/pixel, and acceleration factor *R* = 2) were acquired for anatomical reference and cortical surface reconstruction. *Functional data:* A single-shot gradient-echo EPI sequence, using simultaneous multi-slice (SMS) with blipped Controlled Aliasing in Parallel Imaging (CAIPI) sequence (Setsompop et al., 2012), was used to acquire the GE-BOLD fMRI data at three graded spatiotemporal resolutions. *Acq. 1*: 1.1-mm isotropic nominal voxel size, TR/TE = 1700/26 ms, flip angle = 72°, FOV = 192×192 mm^2^, 87 slices with no gap, acceleration factor *R* = 4, multiband factor = 3, nominal echo spacing = 0.79 ms, bandwidth = 1512 Hz/pixel; *Acq. 2*: 1.5-mm isotropic nominal voxel size, TR/TE = 1120/23 ms, flip angle = 62°, FOV = 192×192 mm^2^, 63 slices with no gap, acceleration factor *R* = 3, multiband factor = 3, nominal echo spacing = 0.69 ms, bandwidth = 1776 Hz/pixel; *Acq. 3*: 2.0-mm isotropic nominal voxel size, TR/TE = 928/26 ms, flip angle = 58°, FOV = 192×192 mm^2^, 51 slices with no gap, acceleration factor *R* = 2, multiband factor = 3, nominal echo spacing = 0.57 ms, bandwidth = 2170 Hz/pixel.

### 3.3 Cortical depth estimation

The high-resolution MPRAGE images were bias corrected (Zaretskaya et al., 2018) and used to automatically reconstruct cortical surfaces with FreeSurfer (Fischl, 2012) (https://surfer.nmr.mgh.harvard.edu). Normalized or “equidistant” cortical depth (‘0%’: white/gray matter boundary; ‘100%’: pial surface) was computed for each EPI voxel using a *voxel-based cortical-depth analysis* under which depth is defined as the relative distance between the voxel centroid coordinate and the nearest white surface and pial surface normalized by the cortical thickness at the corresponding cortical location (Polimeni et al., 2018, 2010). Depths < 0% denote white matter locations below the cortex and depths > 100% denote CSF locations above the cortex (see Fig. 1).

### 3.4 Preprocessing and ICA

Following standard slice-time and motion correction (rigid-body co-registration) using AFNI (Cox, 1996) (https://afni.nimh.nih.gov), functional images were spatially decomposed into multiple independent components (ICs) using FSL’s MELODIC ICA (https://fsl.fmrib.ox.ac.uk/fsl) with the optimal component number estimated using Bayesian dimensionality estimation techniques (Beckmann, 2012; Beckmann and Smith, 2004). Functional images were spatially smoothed until they reached 3D FWHM = 2 mm using AFNI *3dBlurToFWHM* to increase the image SNR for ICA. Of note, this spatial smoothing step was only applied as a preprocessing step for ICA, i.e., to compute ICs; estimations of CortiLag and task activation were performed on the non-smoothed data.

### 3.5 Differentiating between BOLD and non-BOLD components based on CortiLag patterns

After decomposing the 4-D fMRI data into multiple spatial components using ICA, we categorized the resulting ICs as BOLD or non-BOLD components according to the associated CortiLag patterns, which we refer to as “CortiLag-ICA” (as it is akin to assessing TE dependence in ME-ICA), as illustrated in Fig. 2.

**Figure 2:**
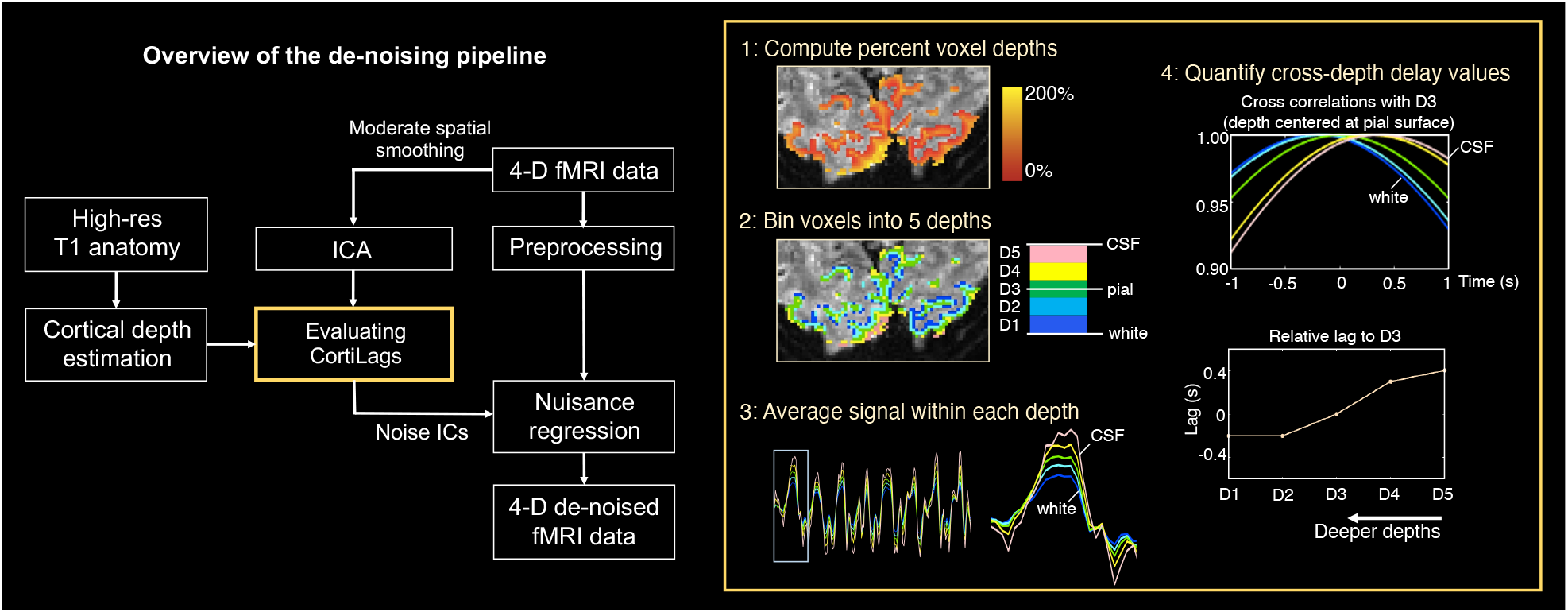
Overview of the proposed CortiLag-ICA de-noising scheme. The across-cortical-depth delay pattern of each IC was characterized following steps highlighted in the yellow box; identified noise components were then included as additional nuisance covariates in the GLM to infer the visual task activation of interest.

For each IC, above-threshold voxels (z-score > 2.3, direct outputs from MELODIC following mixture modeling) were retained and separated into five groups based on their normalized cortical depths (D1: 0–40%, D2: 40–80%, D3: 80–120%, D4: 120–160%, D5: 160– 200%). The representative signal of each cortical depth was derived by averaging the voxel-wise fMRI time series data (after basic slice-time and motion correction) across all voxels within the cortical-depth group. To enable detection of sub-second cross-depth temporal lags, each group’s signal was temporally up-sampled to a 0.1-s grid. The time series of each voxel was weighted by the z-score of the corresponding IC map prior to this averaging. Relative temporal lags between depth-specific signals and the D3 signal were estimated using temporal cross-correlation, yielding a five-element temporal lag vector, including positive and negative lags (with the third element being 0-s lag), that represents the global CortiLag pattern of each IC. We then tested whether the CortiLag pattern was consistent with greater positive lags of responses sampled from the superficial depths, representing a greater time delay, to determine whether an IC was consistent with a proper BOLD signal component. Specifically, we correlated the five-element lag vector of each IC with an ‘order vector’, i.e., a five-element vector of the integers 1 to 5, using Spearman’s rank correlation (i.e., the correlation of ordered ranks of the elements of the two vectors), yielding a coefficient *r*_lag_ (with *r*_lag_ = 1 indicating monotonically increasing temporal lags across depths). A rank correlation metric was used because hemodynamic delays are not expected to be a strictly linear function of cortical depth (Jung et al., 2021). ICs with both high *r*_lag_ values (≥ 0.2) and high *t*_lag_ values (≥ 0.2 s, defined as the temporal lag between D5 and D1 signals) were labeled as BOLD signal components. As shown in Supplementary Fig. S3, among all ICs, *r*_*lag*_ = 0.2 clearly separates high and low rank correlation regimes, and *t*_*lag*_ values exhibited a steeper increase beyond 0.2 s. These thresholds were therefore chosen to define signal and noise components.

### 3.6 Influence on visual task activation

We evaluated the performance of this CortiLag-ICA de-noising framework using visual task data at graded spatiotemporal resolutions. Task activation in response to flickering checkerboard stimuli was inferred using a standard general linear model (GLM) framework, with the task-evoked response modeled by convolving the stimulus paradigm timing with a canonical hemodynamic response function (Glover, 1999). The temporal derivative of the modeled time series was also included as a covariate in the GLM to account for regional delays in the evoked responses.

To demonstrate the ability of our framework to identify and remove noise components, for each resolution the non-BOLD components identified using the procedure described above in *Section 3.5* were first orthogonalized against the stimulus responses and then included as additional nuisance regressors in the GLM analysis.

To benchmark the efficiency of the CortiLag-ICA framework, we also examined an anatomical component-based method (aCompCor) (Behzadi et al., 2007) for noise removal. White-matter and CSF masks were derived from FreeSurfer segmentation, and further eroded using the *3dmask_tool* in AFNI (with erosion steps of 4, 3, and 2 for the 1.1-, 1.5-, and 2.0-mm isotropic data, respectively). The first five principal components of time-series data within the combined white-matter and CSF masks were extracted as the aCompCor nuisance regressors. These aCompCor nuisance regressors, along with six rigid-body motion parameters, were orthogonalized against the stimulus responses and then included as additional nuisance regressors in the GLM analysis to quantify task activation.

For all GLM analyses, serial correlations of the residuals were modelled using the SPM FAST option (https://www.fil.ion.ucl.ac.uk/spm/software/spm12/).

### 3.7 Weighted combinations of fMRI data across cortical depths

Having established the CortiLag-ICA framework of cleaning BOLD-fMRI data, we further tested whether forming suitable weighted combinations of the fMRI data across different depths could result in improved functional sensitivity, in a manner analogous to the optimal across-echo averaging in ME-ICA method. In contrast to the analyses described above where we estimated the normalized cortical depth of each voxel and performed a voxel-based cortical-depth analysis, in the analysis described in this section we project the fMRI data onto each subject’s native surface space and employ a *surface-based cortical-depth analysis*. For this, we first generated four intermediate cortical surfaces equally spaced between the white and pial surfaces (i.e., four equi-distant intracortical surfaces at 20, 40, 60, and 80% cortical depth) between the white matter (0%) and pial (100%) surfaces using FreeSurfer.

This proof-of-principle analysis was carried out using the 1.1-mm iso. visual task data. We projected the 4-D fMRI data onto each surface mesh using the nearest-neighbor interpolation. This analysis assumes that the neuronal activity is similar across cortical depths (i.e., that the functional organization is largely “columnar”) (Polimeni et al., 2018). The combined signal at each vertex of white-matter surface mesh was computed from the weighted average of fMRI fluctuations across corresponding vertices of the white and intracortical surface meshes (i.e., at 0, 20, 40, 60, and 80% cortical depth). Note that supra-millimeter voxels may intersect with more than one cortical surface mesh spaced across depths, thus all voxels intersecting with the pial surface were excluded from this weighted average to mitigate potential contamination from large pial veins. To test whether functional sensitivity (*t*-score and the spatial extent of task activation) can be boosted using a plausibly physiologically-informed weighted combination, signals of each cortical depth were first converted to percent signal changes (PSCs) and then weighted by the depth-specific PSC level predicted by a cortical vascular model: the fMRI data from cortical depths [0% 20% 40% 60% 80%] were weighted by [0.014 0.022 0.029 0.032 0.037] (i.e., the modeled depth-specific PSC levels in Fig. 3b from (Markuerkiaga et al., 2016)), respectively, prior to combination. This model-based PSC weighting scheme is termed as “mPSC”, analogous to SNR-weighted combination under the assumption of uniform noise levels across depths. As a control analysis, we also tested two alternative approaches to combining the fMRI data across cortical depths: (*i*) assigning equal weights to all depths (‘equal weights); and (*ii*) weighting each depth by the reverse order of mPSC, i.e., [0.037 0.032 0.029 0.022 0.014] (‘rev mPSC’). We then conducted GLM analyses on the combined across-depth time-series data using different weighting schemes to assess visual task activation. Statistical scores resulting from the two control analyses were expected to be smaller than those derived using the proposed “mPSC” combination.

**Figure 3:**
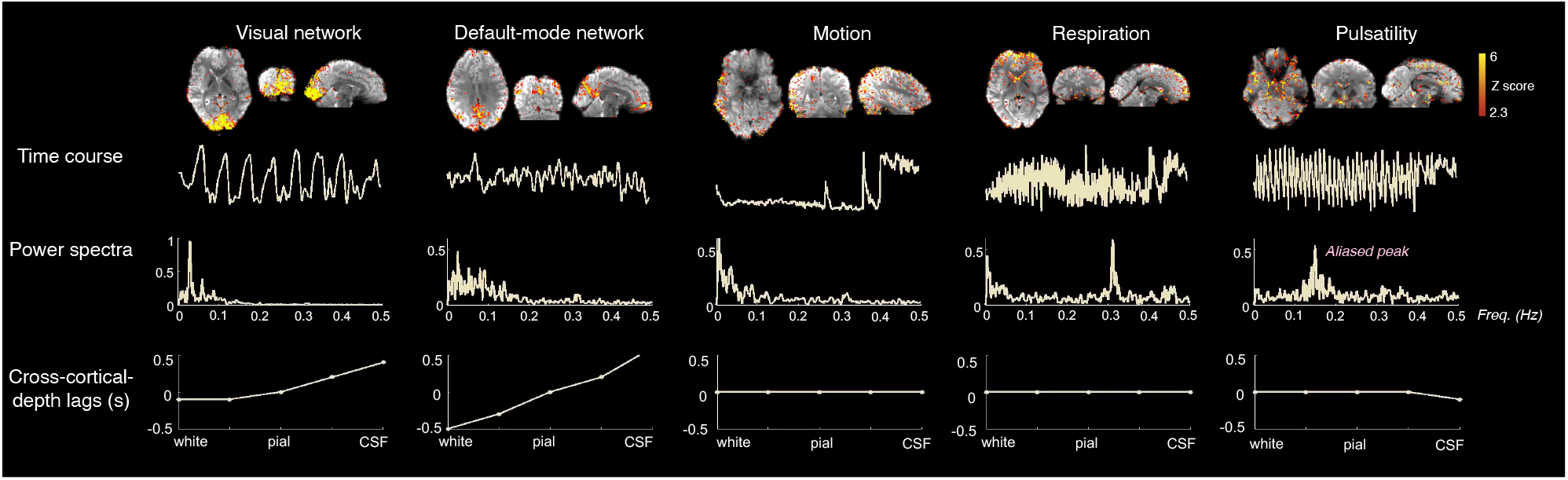
Distinct functional BOLD signal components (“Visual network” and “Default-mode network” ICs) and non-BOLD noise components (subject “Motion”, “Respiration” and “Pulsatility” ICs) identified using the CortiLag patterns, illustrated using a 2-mm iso. visual task data from a representative subject. Top row: the spatial map of each example IC; middle rows: the time course and the power spectrum of the time course corresponding to each IC; bottom rows: the corresponding temporal lag values of each cortical depth (D1–D5) relative to the middle depth (D3). Note that the cardiac peak was aliased into low frequencies (∼0.15 Hz) with this TR (0.928 s) used for this acquisition (“Pulsatility” IC); see Supplementary Fig. S2 for another illustrative case in which the cardiac peak was resolved at a shorter TR value.

## 4 Results

### 4.1 Distinct CortiLag patterns of functional BOLD signal components and non-BOLD noise

Figure 3 illustrates that distinct CortiLag patterns reflecting either functional signal components or noise components can be identified in the measured GE-BOLD fMRI data. In this representative dataset, the functional connectivity patterns that correspond to resting-state networks (“Visual network” and “Default-mode network”) exhibited the expected progression of temporal delays from deep to superficial depths; apparent artifacts, including motion and quasi-periodical fluctuations time-locked to respiratory and cardiac cycles, exhibited no clear delays across cortical depths.

Overall, as illustrated by the IC sorting results of an exemplar subject (Fig. 4), ICs with the highest *r*_lag_ and *t*_lag_ values, i.e., ICs exhibiting a delay from deeper to superficial depths, generally exhibited structured spatial patterns that resembled known task-relevant or resting-state functional networks; ICs with zero across-depth temporal lags were associated more with physiological (> 0.1-Hz fluctuations) or motion-like spatial patterns (high z-scores at the edges of the brain), suggesting that the CortiLag-ICA framework can isolate functional BOLD signal components from common forms of non-BOLD noise at the spatiotemporal resolutions of our dataset, as intended. IC statistics and the distributions of *r*_lag_ and *t*_lag_ values across all ICs of each subject are summarized in Supplementary Fig. S3.

**Figure 4:**
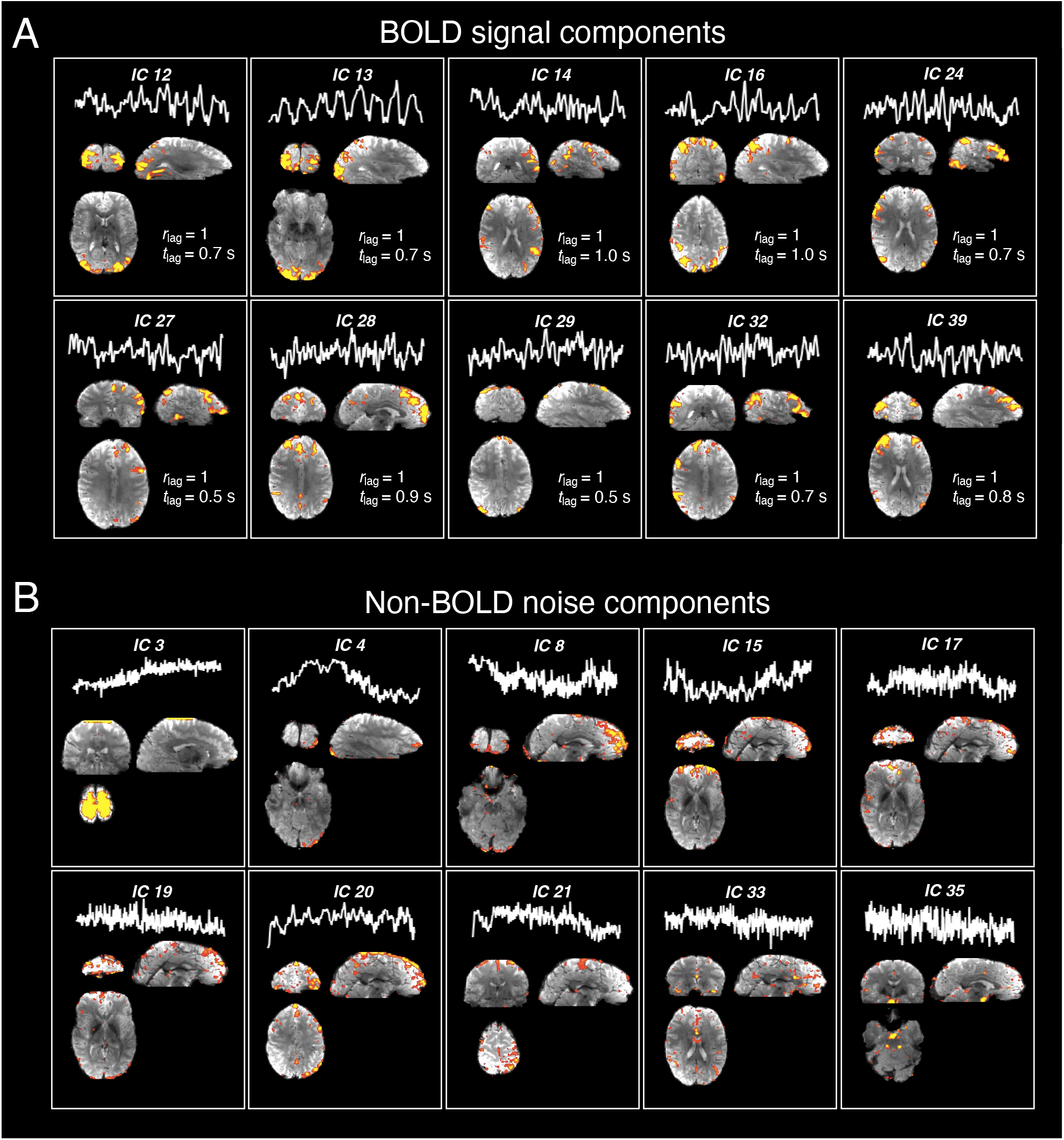
Illustration of the efficacy of employing CortiLag patterns to differentiate between BOLD signal and non-BOLD noise components. (A) Exemplar functional BOLD signals (ICs with high *r*_lag_ and *t*_lag_ values) and (B) non-BOLD noise (ICs with low *r*_lag_ or *t*_lag_ values; all displayed ICs showed zero across-depth lags), demonstrated using the 1.5-mm iso. visual task data from a representative subject. Each panel shows the time course and spatial pattern of each IC. Components with the highest *r*_lag_ and *t*_lag_ values comprised the task-active (IC 13) and common functional networks; whereas components with the lowest *r*_lag_ or *t*_lag_ values resembled typical structured-noise patterns or artifacts.

### 4.2 Enhanced sensitivity to visual activation post removal of non-BOLD ICs

Figures 5 and 6 present individual- and group-level visual task activation results for the minimally processed data following basic rigid-body co-registration and slice-time correction (“raw”), data denoised by including the aCompCor plus motion regressors (“aCompCor”), and the data denoised by including the non-BOLD ICs (*r*_lag_ < 0.2 or *t*_lag_ < 0.2 s) as nuisance covariates in the GLM analysis (“CortiLag”). Across all resolutions, CortiLag regressors led to significantly greater reductions in total residual variance in the GLM analyses—averaging 16%, 10%, and 8% in primary V1 for the 2.0-, 1.5-, and 1.1-mm data, respectively—compared to aCompcor regressors, which achieved average reductions of 12%, 6%, and 5% for the same resolutions, after accounting for differences in degrees of freedom. This suggests that the nuisance components identified by the CortiLag criteria more effectively capture the regional physiological and motion artifacts in gray matter than those derived from white matter and ventricles and those based on rigid-body motion parameter estimates. Accordingly, enhanced task sensitivity was observed after de-noising at all imaging resolutions. Relative to the “raw” condition, CortiLag-ICA increased the mean *t*-score in V1 by 16%, 9%, and 9% for the 2.0-, 1.5-, and 1.1-mm data, respectively. In contrast, aCompCor yielded increases of 14%, 4%, and 5% for the same resolutions. The *t*-scores were directly compared between different de-noising framework, as their associated *t*-distributions approximated a normal distribution due to the large degrees of freedom (well over 30), despite differences in the number of nuisance regressors identified in each method. These results demonstrate the efficacy of the CortiLag-ICA framework in de-noising moderate- and high-resolution fMRI data.

**Figure 5:**
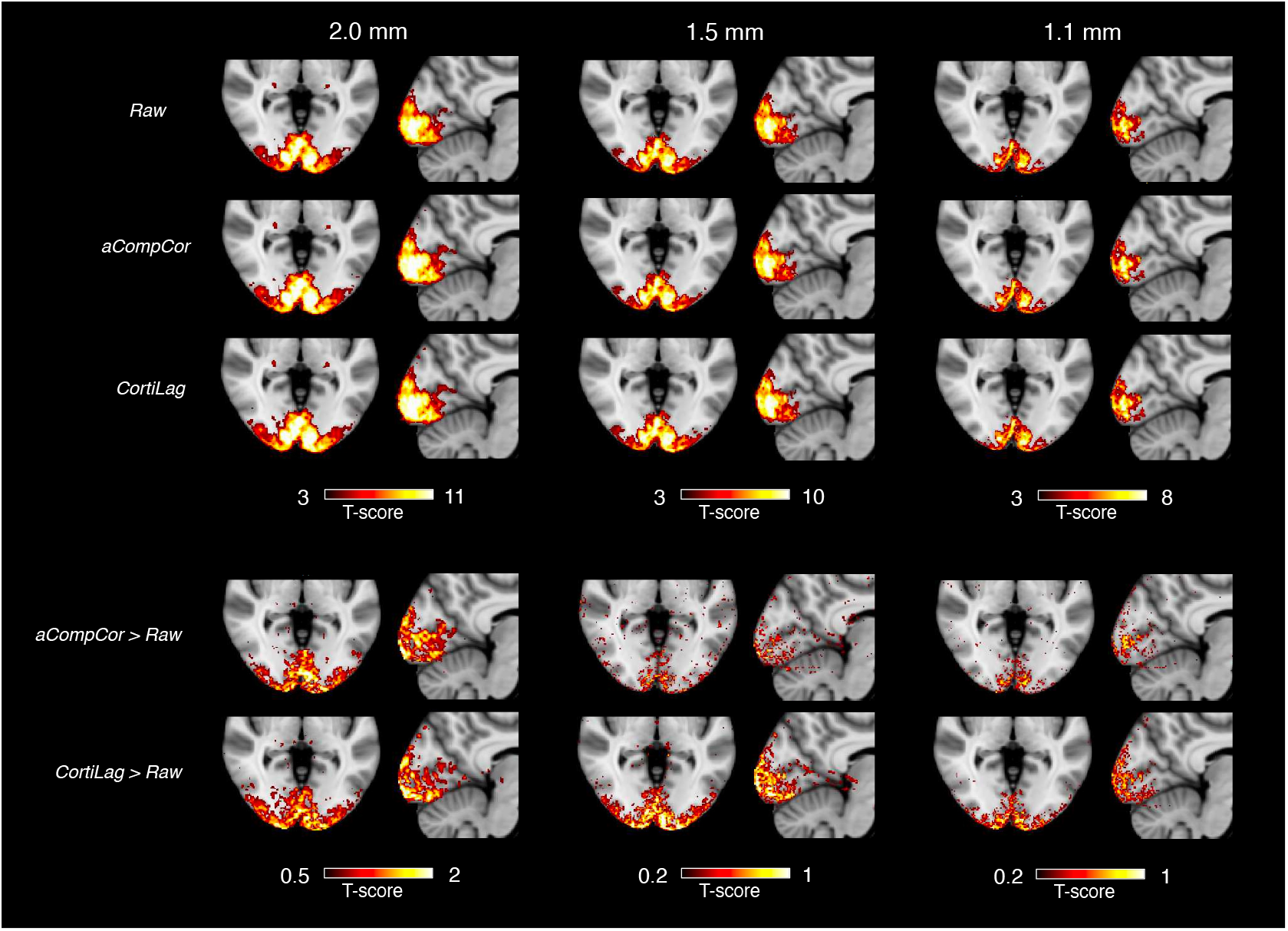
Group-mean visual task activation (N=12). “raw”: fMRI data after basic preprocessing (rigid-body co-registration and slice-time correction); “aCompCor”: including six motion parameters and five aCompCor principal components as nuisance regressors in the GLM analysis; “CortiLag”: including noise ICs (*r*_lag_ < 0.2 or *t*_lag_ < 0.2 s) as nuisance regressors in the GLM analysis. To facilitate visual comparison of activation results across preprocessing pipelines, difference maps are shown at the bottom: “aCompCor > Raw” shows the difference between the “aCompCor” and “Raw” *t*-score maps, and “CortiLag > Raw” represents the difference between “CortiLag” and “Raw” *t*-score maps. Individual-level *t*-score results were normalized to the MNI152 template using ANTs (Avants et al., 2011).

**Figure 6:**
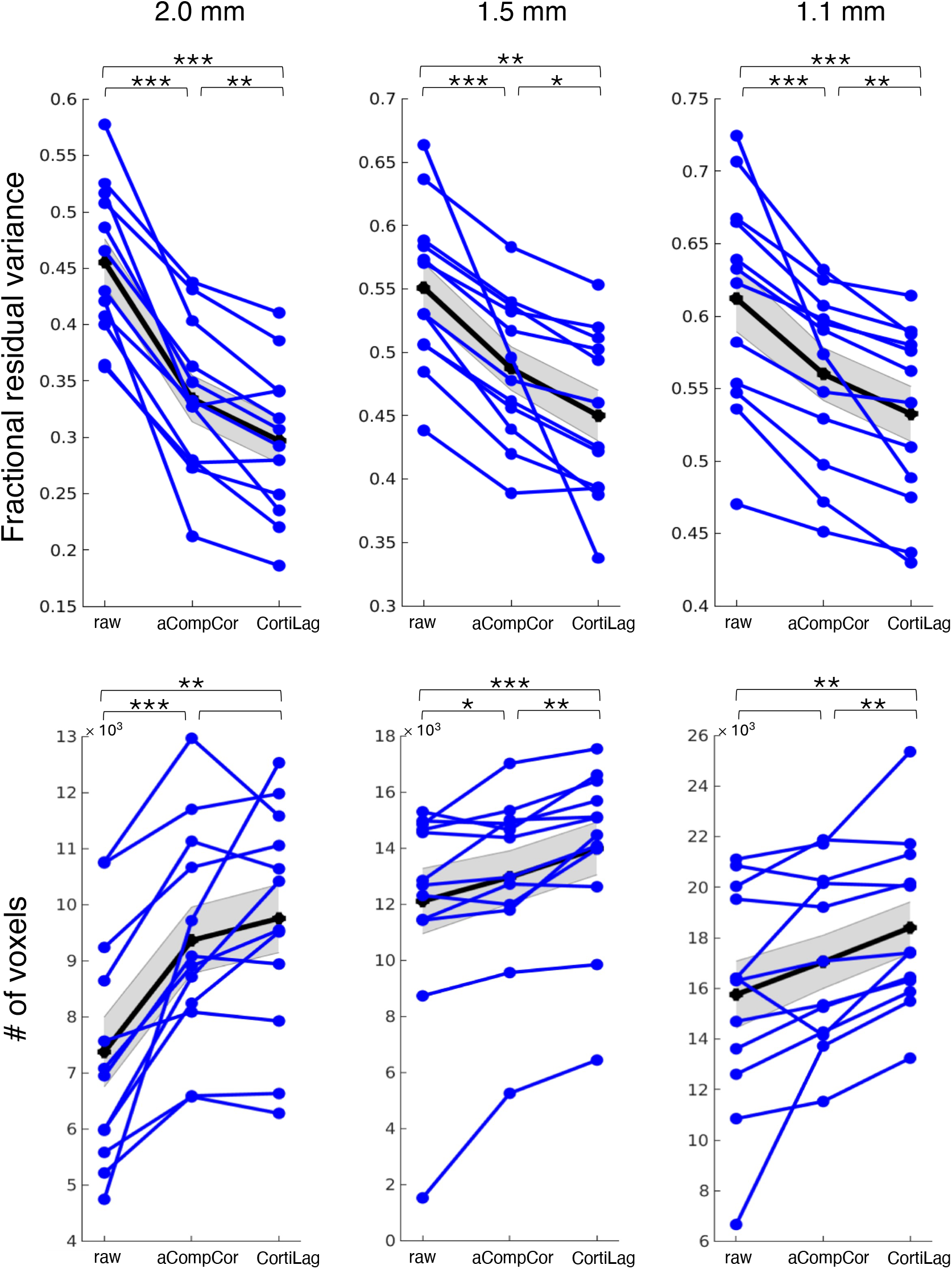
Influence of de-noising on visual task activation. Both (A) the mean squared error of residual variance in GLM analyses (fraction relative to the total variance of each voxel’s time series, corrected for changes in degrees of freedom) within the FreeSurfer segmented primary visual cortex, V1 (“Fractional residual variance”) and (B) task-active voxel counts (“# of voxels”, *t*-score > 3, FDR < 0.05) in the entire occipital cortex were quantified for comparison. “raw”: fMRI data after basic preprocessing (rigid-body co-registration and slice-time correction); “aCompCor”: including six motion parameters and five aCompCor principal components as nuisance regressors in the GLM analysis; “CortiLag”: including noise ICs (*r*_lag_ < 0.2 or *t*_lag_ < 0.2 s) as nuisance regressors in the GLM analysis. Each blue line represents the results of a single subject. Mean and standard errors across subjects are displayed in gray. Statistical significance of paired *t*-test results (“aCompCor” vs. “CortiLag”) is indicated above the columns of de-noising summary metrics: (*) *p* < 0.05; (**) *p* < 0.005; (***) *p* < 0.0005.

### 4.3 Optimal combination of information across cortical depths

Having established that the CortiLag-ICA framework can be employed to de-noise GE-BOLD fMRI data, we additionally evaluated whether weighted combination of information across all cortical depths could further enhance the sensitivity to detect brain activity, compared to the activation estimated using a single depth. Figure 7 summarizes the statistical estimates of visual task activation at each cortical depth and following different schemes of weighted combinations of the data across depths. It is noteworthy that, at the group level, weighting the data sampled at each depth by unoptimized, model-based PSCs (“mPSC”) derived from an independent simulation already yielded higher functional sensitivity (largest number of statistically-significant vertices) than the activation measured at any given depth, as well as the other two schemes shown for comparison (i.e., weights derived from uniform values (“equal weights”), or the reverse of mPSCs (“rev mPSC”)). These observations suggest that an informed weighted combination of data across cortical depths could further enhance the functional sensitivity of small-voxel GE-BOLD fMRI data.

**Figure 7:**
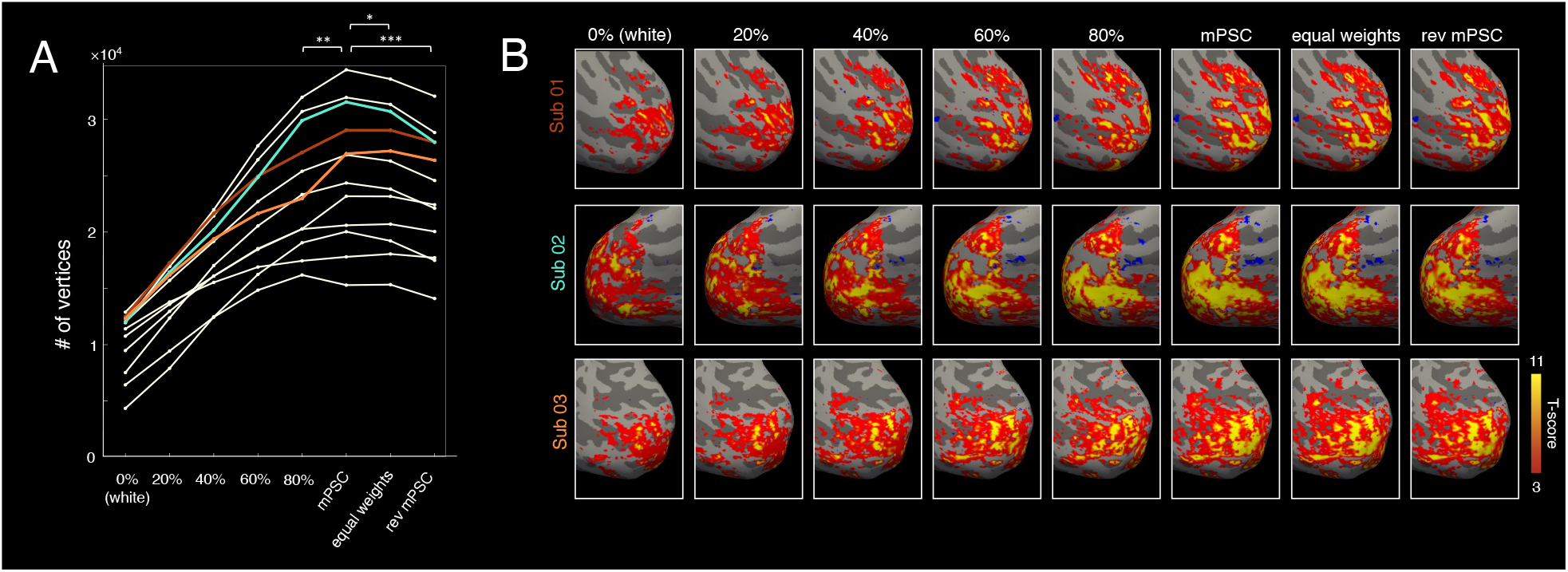
Impact of different schemes of weighted combination of fMRI data across cortical depths on functional sensitivity, illustrated using the 1.1-mm iso. visual task data. (A) Task-active (*t*-score > 3 and FDR < 0.05) vertex counts at different cortical depths and after weighted combination of data across all depths (“mPSC”: weights given by the modeled PSC of each depth (Markuerkiaga et al., 2016); “equal weights”: uniform weights across all depths; “rev mPSC”: weights given by the reverse order of modeled PSC across depths). Each trace represents results of a single subject. Statistical significance of paired *t*-test results (“mPSC” vs. “80%”, “equal weights”, and “rev mPSC”) is indicated above the associated pairs of weighting schemes: (*) *p* < 0.05; (**) *p* < 0.005; (***) *p* < 0.0005. (B) Visual task activation maps presented on the inflated cortical surface for three representative subjects, thresholded to facilitate visual comparison across different combination methods and to illustrate improved statistical scores relative to cortical depth-specific results.

## 5 Discussion

### 5.1 General findings

In this work, we demonstrated the feasibility of leveraging cortical-depth-dependent analysis to enhance the sensitivity and neuronal specificity of moderate- and high-spatial-resolution GE-BOLD fMRI data. First, we showed that functional BOLD signal components and non-BOLD noise components exhibited distinct CortiLag patterns, which could aid in identifying and removing noise fluctuations from fMRI data. We then demonstrated that, by appropriately combining data across cortical depths, the overall statistical sensitivity to detect functional activity could be further increased.

### 5.2 Comparison with alternative data-driven approaches

The CortiLag-ICA de-noising framework proposed here is analogous in many ways to ME-ICA. However, instead of evaluating the dependence on TE, we mapped the temporal progression of hemodynamic changes across cortical depths to distinguish BOLD signal components from potential noise components. Additionally, rather than forming weighted combinations of the fMRI time-series data across echoes, we formed weighted combinations of fMRI data across cortical depths to improve CNR.

In theory, the CortiLag-ICA framework should achieve denoising efficacy comparable to that of ME-ICA for GE-BOLD fMRI. Typical nuisance factors that exhibited no TE dependence— e.g., intensity changes related to head movements, quasi-periodical respiration, pulsatility and imaging acquisition/reconstruction artifacts—are not expected to exhibit graded, hundred milliseconds-scale temporal lags across cortical depths, and thus ought to be categorized as noise under the CortiLag framework as well. Previous studies have cautioned that TE dependence alone cannot separate components of interest from nuisance components, since many forms of physiological noise will generate changes in blood oxygenation and thus will manifest as BOLD-like components, while neuronal activation may cause non-BOLD modulations. For example, slow hemodynamic variations driven by the autonomic nervous system (i.e., changes in heart rates and end-tidal CO_2_ levels (Birn et al., 2008; Shmueli et al., 2007; Wise et al., 2004)) will cause BOLD-like changes similar to changes driven by local neuronal activity (Kundu et al., 2012; Power et al., 2018), given that both dynamics generate changes in blood oxygenation and therefore will exhibit “BOLD-like” signal changes in T_2_*-weighted acquisitions. Also, neuronal activation can cause inflow effects, blood volume changes, and tissue/CSF displacement, which may generate non-BOLD intensity modulations that appear in T_2_*-weighted data (Gao et al., 1996; Jin and Kim, 2010; Liu et al., 2008; Piechnik et al., 2009; Scouten and Constable, 2008), especially when using short TR values that impart additional T_1_ weighting. Unfortunately, this limitation extends to the CortiLag-ICA method as well. Although these “systemic” changes in blood oxygenation propagate through the cortical vasculature in a purely anterograde fashion, as opposed to expected “neurogenic” changes triggered by neurons that cause retrograde dilations progressing from parenchymal microvessels to the feeding arterioles, the neurogenic hemodynamics will also result in anterograde draining of deoxygenated hemoglobin into the veins, and so in both cases we would expect that the corresponding BOLD changes would appear first in the parenchymal and then later in time in pial veins. Supplementary Fig. S4 offers an illustrative case in which an IC correlated with respiratory variation also demonstrates apparent timing differences across depths. Collectively, CortiLag- and ME-ICA are similarly limited in their ability to separate BOLD-like noise components from those signals of interest.

Apart from sharing strengths and limitations with the ME-ICA framework, a major advantage of the CortiLag-based framework is that it can be applied to single-echo acquisitions. Thus our framework may be preferable for fMRI applications that necessitate high spatiotemporal resolutions, e.g., recent studies into rapid brain oscillations (Lee et al., 2013; Lewis et al., 2016), and imaging of small nuclei in brainstem regions (Bianciardi et al., 2016; Sclocco et al., 2018). Yet the CortiLag-based framework also faces additional limitations. First, to resolve timing differences across cortical depths, both the spatial and temporal resolution of fMRI must fall above certain thresholds. According to our results, the graded delays across cortical depths can be well detected in data with TR values up to 1.7 s, and voxel sizes up to 2 mm. This suggests that our method may be compatible with existing large-scale neuroimaging datasets such as that provided by the Human Connectome Project (Van Essen et al., 2013). Second, since the CortiLag characterization relies on the stereotyped pattern of blood supply and drainage to the cortex, it may have diminished denoising efficiency in acquisitions with limited cortical coverage. Third, while the CortiLag method is theoretically feasible even with imperfect cortical depth delineation and surface reconstruction—as long as inferior and superior depths are sufficiently separated to detect lags in the BOLD-fMRI data—its dependence on the accuracy of cortical depth estimation warrants further evaluation in future studies.

Of note, while we present CortiLag-ICA as an alternative to ME-ICA throughout this article, it can also be readily combined with ME-ICA to enhance the overall statistical sensitivity of BOLD fMRI. Although echo- and cortical-depth-dependent information may appear redundant in the denoising step, integrating information across echoes and cortical depths jointly can be complementary, i.e., weighted combinations across cortical depths can be performed in addition to BOLD-sensitivity-weighted combinations across acquired echoes to maximize the overall functional sensitivity of a dataset.

In addition to ME-ICA, approaches based on pretrained classifiers have also gained popularity, such as FMRIB’s ICA-based Xnoiseifier (FIX) (Griffanti et al., 2014; Salimi-Khorshidi et al., 2014) and ICA-based strategies for Automatic Removal of Motion Artifacts (AROMA) (Pruim et al., 2015). However, the accuracy of these methods depends on the availability of suitable training datasets. A preliminary assessment of FIX applied to our moderate-resolution fMRI data (1.5-mm and 2.0-mm isotropic voxel sizes), using a classifier trained on the public 7T HCP dataset (Thanh Vu et al., 2017), demonstrated general consistency in noise component classification with our proposed CortiLag-ICA approach (see Supplementary Methods I and Figs. S5 and S6). This suggests that the CortiLag-ICA method could be a promising alternative for fMRI de-noising, particularly in the absence of either multi-echo acquisitions and suitable training datasets.

### 5.3 Outlook

#### 5.3.1 Applicability to higher- and lower-resolution fMRI data

Theoretically, higher spatial resolution (e.g., sub-millimeter voxel sizes acquired on ultra-high field scanners) allows timing differences across depths to be better resolved for more precise detection of depth-specific vascular signals, thereby improving the accuracy of signal and noise differentiation using CortiLag-ICA. However, in practice, increased spatial resolution often comes at the cost of reduced temporal resolution—hence as voxel size decreases, the limiting factor for CortiLag-ICA performance may become the ability to resolve depth-dependent fluctuations in time. In this study, we tested temporal resolutions up to 1.7 s, future work may explore the feasibility of resolving temporal lags at even lower temporal resolutions. Additionally, with improved spatial specificity and finer divisions across cortical depths (>5 depth divisions as performed in this study), the characterized neurogenic across-depth lag patterns may deviate from a simple monotonic increase, and instead exhibit, e.g., a flat onset followed by a ramp, particularly in scenarios where neuronal activity originates in mid-cortical depths. In such cases, the quantitative criteria for signal and noise identification may need to be adapted to better capture these neurogenic signal features.

Then what is the lowest spatial resolution suitable for CortiLag-ICA? Here, we only reported denoising results using voxel sizes up to 2-mm isotropic spatial resolution. However, theoretically, due to the “extravascular blooming” effect, the physical distance between the white matter surface (having the earliest hemodynamic responses) and the CSF regions (having the latest detectable responses) with measurable BOLD signals can range from 2–6 mm. This implies that the proposed denoising approach may be applicable to acquisitions at even lower spatial resolutions. Additionally, even in locations where the cortex is thin such that its thickness is similar to the linear dimension of the voxel size (such as the primary visual cortex, which in humans is ∼1.5 to 2.5 mm in thickness), given that each voxel samples a distinct depth of the folded cortex, pooling across all voxels within an IC spanning multiple cortical areas, or even within a single cortical area, effectively increases the number of distinct cortical depth sampled (Polimeni et al., 2018). As a preliminary test of this idea, we evaluated the performance of CortiLag-ICA on five additional datasets with 3-mm isotropic voxel sizes (using data from (Blazejewska et al., 2019)) and enhanced task activation was observed after removing noise ICs from the lower-resolution data (see Supplementary Methods II and Supplementary Fig. S7 for detailed descriptions of the relevant data and results). Thus the proposed framework shows a potential to aid de-noising fMRI data acquired at more conventional spatial resolutions when neither external physiological recordings nor multi-echo acquisitions are available. It is of note that as voxel size increases, the absolute across-depth lag differences are expected to gradually diminish, potentially requiring a reassessment of the thresholds (in particular, the threshold used for *t*_*lag*_) used to distinguish signal from noise.

An additional related factor not directly examined in this study is the effect of spatial smoothing— a common preprocessing step used to improve SNR in high-resolution or layer-specific fMRI by averaging uncorrelated noise (Blazejewska et al., 2019; Huber et al., 2021). Due to its complementary nature with CortiLag-ICA (a form of nuisance regression), which enhances the neuronal specificity of the fMRI time series by projecting out noisy fluctuations including structured noise sources, these steps can be combined. Conventional volumetric 3D smoothing kernels blur signals across cortical depths and are thus best applied after CortiLag-ICA. In contrast, local anatomically-informed smoothing along the cortical depth direction is expected to have a smaller effect on de-noising performance, and its application before or after CortiLag-ICA is likely less critical. Future studies could specifically evaluate how distinct types of spatial smoothing interact with CortiLag-ICA denoising to optimize performance.

#### 5.3.2 “Optimal” integration of information across depths

Unlike large-voxel acquisitions where all cortical depths inherently have equal weights, small-voxel acquisitions permit the flexibility of weighting the depth-specific fractional contributions such that the sensitivity and neuronal specificity of fMRI measures can be balanced or maximized (Blazejewska et al., 2019). In the proof-of-principle analysis using our 1.1-mm iso. visual task data, we demonstrated that weighting depth-specific information by a simple vascular model-based PSC can boost the overall sensitivity to detect task activation in cases where the neuronal activation is known to be similar across depths at any given cortical location. Nevertheless, we do not believe that this particular weighting scheme is the only weighting scheme (or an optimal weighting scheme) that would provide such a boost in sensitivity, however this scheme did provide a useful proof of concept. In practice, rather than using this particular physiologically-motived vascular model, other similar models may also help boost sensitivity, and appropriate region-specific weighting schemes may be derived from empirical measurements from an independent resting-state scan akin to a “functional localizer”. Additionally, one dataset may be collected to serve multiple investigations—for instance, activation elicited by several distinct stimuli or tasks using mixed presentation designs, or functional connectivity estimated with respect to distinct network seeds—and the weights for the cortical-depth combination can therefore be tailored separately to optimize different metrics of interest. This new data-driven and “anatomically-informed” (or perhaps “physiologically-informed”) analysis approach further motivates small-voxel acquisitions and investments in improving imaging resolution (Blazejewska et al., 2019).

## 6 Conclusions

In this work, we demonstrate that the temporal progression of hemodynamic signals across cortical depths can be leveraged to differentiate between BOLD and non-BOLD components in T_2_*-weighted fMRI, effectively enhancing its sensitivity and neuronal specificity. The approach is particularly suitable for moderate- and high-resolution acquisitions when multi-echo acquisitions are less advantageous. Our study also highlights the potential of small-voxel acquisitions in facilitating novel preprocessing and analytical strategies for fMRI, extending beyond their well-established usefulness for producing fine-grained depictions of brain functional architecture.

## Acknowledgements

This work was supported in part by the National Institutes of Health (NIH) including the National Institute of Biomedical Imaging and Bioengineering (grant numbers P41-EB015896, P41-EB030006, R01-EB019437, R01-EB032746, and R01-EB035560), the National Institute on Aging (grant number RF1-AG074008) and the National Institute of Neurological Disorders and Stroke (grant number K99/R00-NS118120 and U19-NS123717), by the National Institute of Mental Health (grant numbers R01-MH111438, R01-MH111419, R21-MH135201, and R00-MH120054), and by the MGH/HST Athinoula A. Martinos Center for Biomedical Imaging; and was made possible by the resources provided by NIH Shared Instrumentation grants S10-RR023043 and S10-RR019371.

