## Supplementary Materials for "Differentiating BOLD and non-BOLD signals in fMRI time series using cross-cortical depth delay patterns"

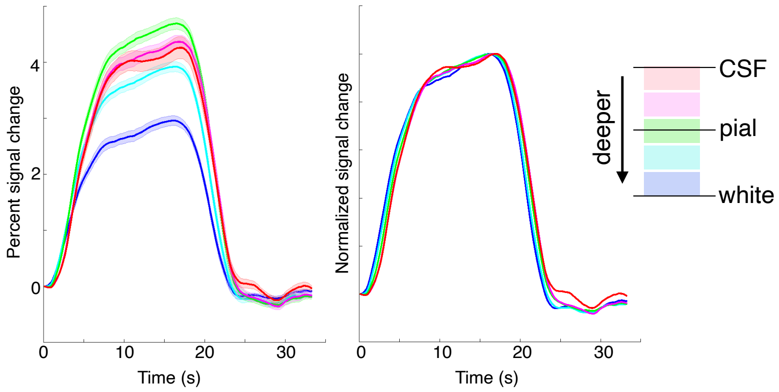


**Figure S1:** Neurogenic hemodynamic responses with graded temporal delays across cortical depths, illustrated using the 1.1-mm visual task data (see Methods). Left: mean responses and standard errors across block trials (stimulus on/off = 16/19 s, *N*=12 subject, 8 blocks per subject); Right: mean responses normalized by peak signal intensity.

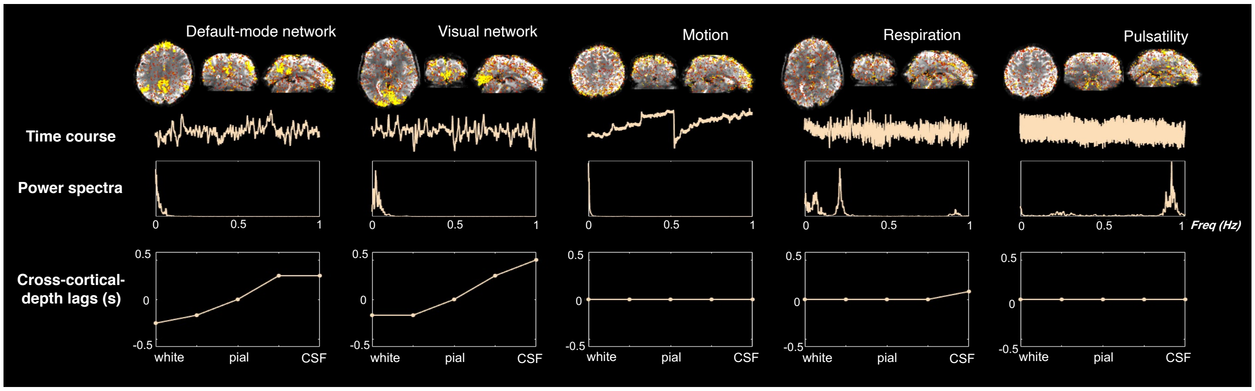


**Figure S2:** Distinct CortiLag patterns observed between BOLD signal components (“Default-mode network” and “Visual network” ICs) and non-BOLD noise components (subject “Motion”, “Respiration” and “Pulsatility” ICs). Top: the spatial map of each IC; middle: the corresponding time course and power spectrum of the time course for each IC; bottom: the temporal lag values of each cortical depth (D1–D5) relative to the middle depth (D3). For this illustrative resting-state dataset, TR values were shortened to resolve the cardiac peak at ~1.0 Hz (2-mm isotropic voxel size, TR = 0.481 s, acceleration factor *R* = 2, multiband factor = 5).

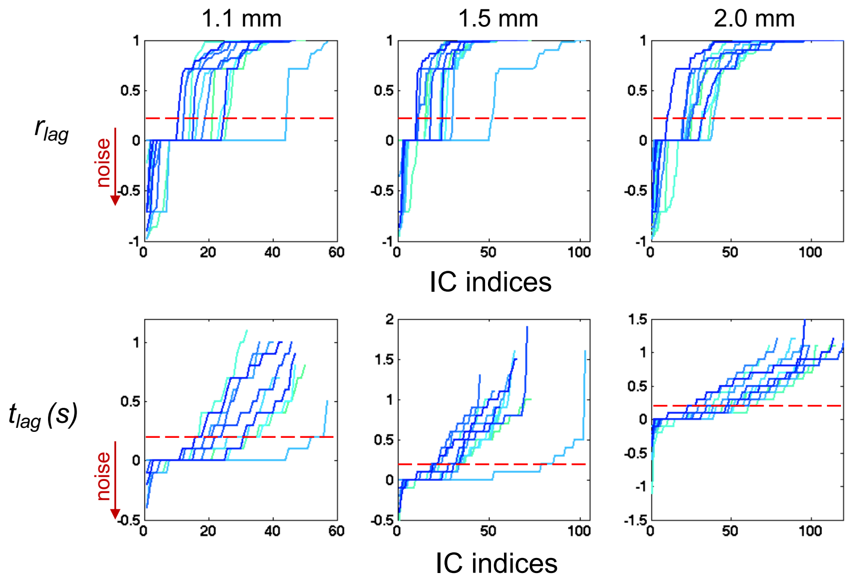


**Figure S3:** Plots of $r_{lag}$ and $t_{lag}$ values across all independent components (ICs) for each subject. Each trace represents an individual participant, progressing from low to high $r_{lag}$ or $t_{lag}$ values across all ICs (*N*=12 subjects, with distinct colors representing the results of each subject). The percentages of noise ICs ($r_{lag}$ < 0.2 or $t_{lag}$ < 0.2 s) were 57±14% (among 43±7 total ICs) for the 1.1-mm iso. data; 49±12% (among 63±15 total ICs) for the 1.5-mm iso. data; and 43±11% (among 98±14 total ICs) for the 2.0-mm iso. data.

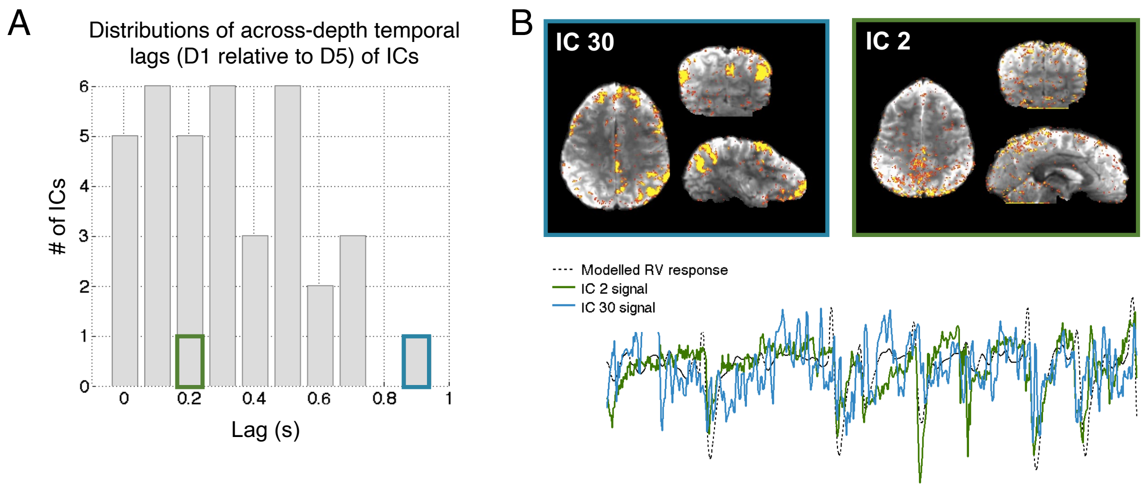


**Figure S4:** Respiratory variation (RV)-correlated ICs, depending on the specific origin, may exhibit either clear or ambiguous temporal lags across cortical depths. For example, panel (A) shows the distributions of temporal lags between cortical depths D1 and D5 (D1 preceding D5) across all ICs in a representative subject. We could identify two ICs correlated with RV: one exhibiting a clear BOLD-like across-cortical-depth temporal lag (blue box), and one that exhibited less discernable timing across cortical depths (green box). Time courses and spatial patterns of these two ICs are shown in panel (B). (The "modelled RV response" was derived from external respiratory recordings following Chang *et al.*[1].)

### Supplementary Methods I: CortiLag-ICA vs. FIX

In addition to comparing task de-noising performance with aCompCor (Figs. 5 and 6 in the main text), we also assessed the consistency of signal IC and noise IC identification between CortiLag-ICA and a widely used ICA-based denoising method, FIX (or "FMRIB's ICA-based Xnoiseifier")[2]. FIX automatically labels signal and noise components based on a pre-trained, hand-labeled dataset. Its performance therefore depends on the similarity between the spatiotemporal characteristics of the fMRI training dataset and the acquisitions of interest. Note that since no existing hand-labeled training dataset matches the specific spatiotemporal properties of our 7T BOLD fMRI acquisitions, the IC classification results provided by FIX cannot be considered ground truth. Nevertheless, this assessment provides valuable insight into the distribution of CortiLag patterns among ICs and the feasibility of using CortiLag-ICA to effectively classify signal and noise components. Furthermore, this assessment provides a helpful, practical comparison between the classification produced by proposed CortiLag-ICA approach and a commonly-used classification produced by FIX, both of which are applied to the same ICA outputs.

Specifically, we used an existing classifier derived from the 7T HCP dataset ("*HCP7T_hp2000.RData*") provided by the FIX toolbox and applied it to our visual task fMRI datasets with 1.5-mm and 2.0-mm isotropic resolutions. We then compared the CortiLag patterns of ICs classified as signal and noise by FIX. We defined the FIX IC probability above 0.5 as "signal" and below 0.2 as "noise". The prediction was that ICs classified by FIX as "signal" should exhibit a clear increase in temporal lag across cortical depths, and ICs classified by FIX as "noise" should exhibit no difference in temporal lag across depths. As expected, CortiLag patterns of ICs classified as signal and noise by FIX showed substantial differences (Supplementary Fig. S5). Indeed, for all subjects, we observed that the average cortical lag pattern for "signal" components exhibited an increase in lag from the white to the pial surface, and the average cortical lag pattern for "signal" components exhibited no discernable change in lag from the white to the pial surface. For individual ICs, at the group level the fraction of FIX-classified signal components also identified as "signal" by CortiLag-ICA was 84 ± 3% (mean and standard deviations across subjects) for the 1.5-mm data and 88 ± 8% for the 2.0-mm data. Similarly, the fraction of FIX-ICA-classified noise components also classified as "noise" by CortiLag-ICA was 84 ± 18% for the 1.5-mm data and 75 ± 13% for the 2.0-mm data.

It is challenging to determine the superiority of one method over the other based on our results, as both rely on heuristic selection of parameter values (e.g., cut-off thresholds for noise identification) and, in the FIX case, the choice of an appropriate training dataset. To demonstrate that each method has similar shortcomings, we further visually inspected ICs with classification results that disagreed between the two methods, and observed that both methods misclassify clear noise components as signal components, with two such examples shown in Supplementary Fig. S6. Nonetheless, our assessment revealed general consistency of noise classification across the two classification methods, supporting the feasibility and efficacy of employing CortiLag patterns to identify and remove noisy ICs—particularly in applications where multi-echo acquisitions or suitable training datasets are unavailable.

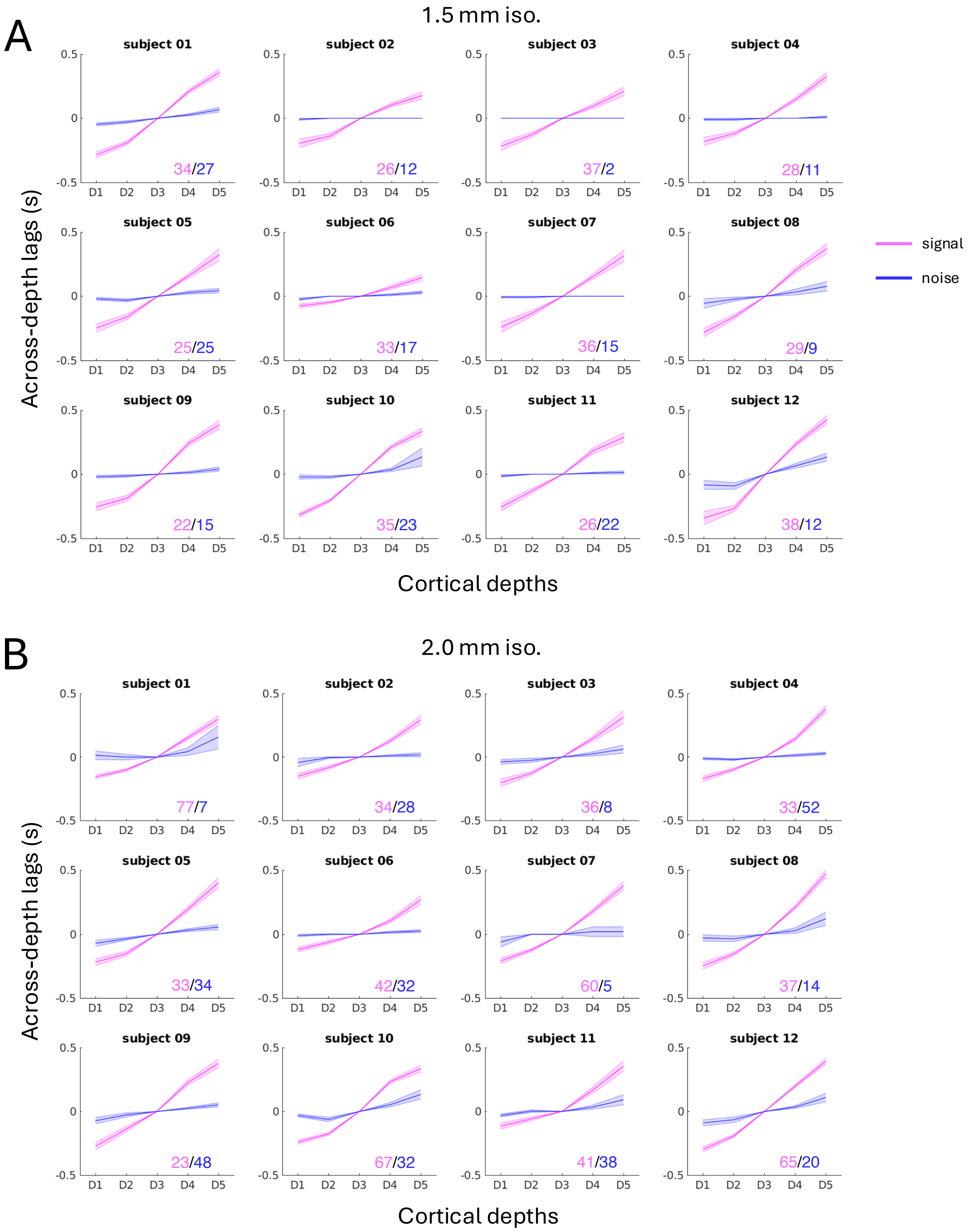


**Figure S5:** CortiLag patterns of "signal" and "noise" components identified by FIX-ICA at two different spatial resolutions (Top: 1.5-mm isotropic voxel size; Bottom: 2.0-mm isotropic voxel size). Mean and standard error (represented by shading) of CortiLag patterns relative to the mid-cortical depth (D3) are displayed for different categories of ICs. The numbers of "signal" and "noise" components are indicated at the bottom-right corner of each panel.

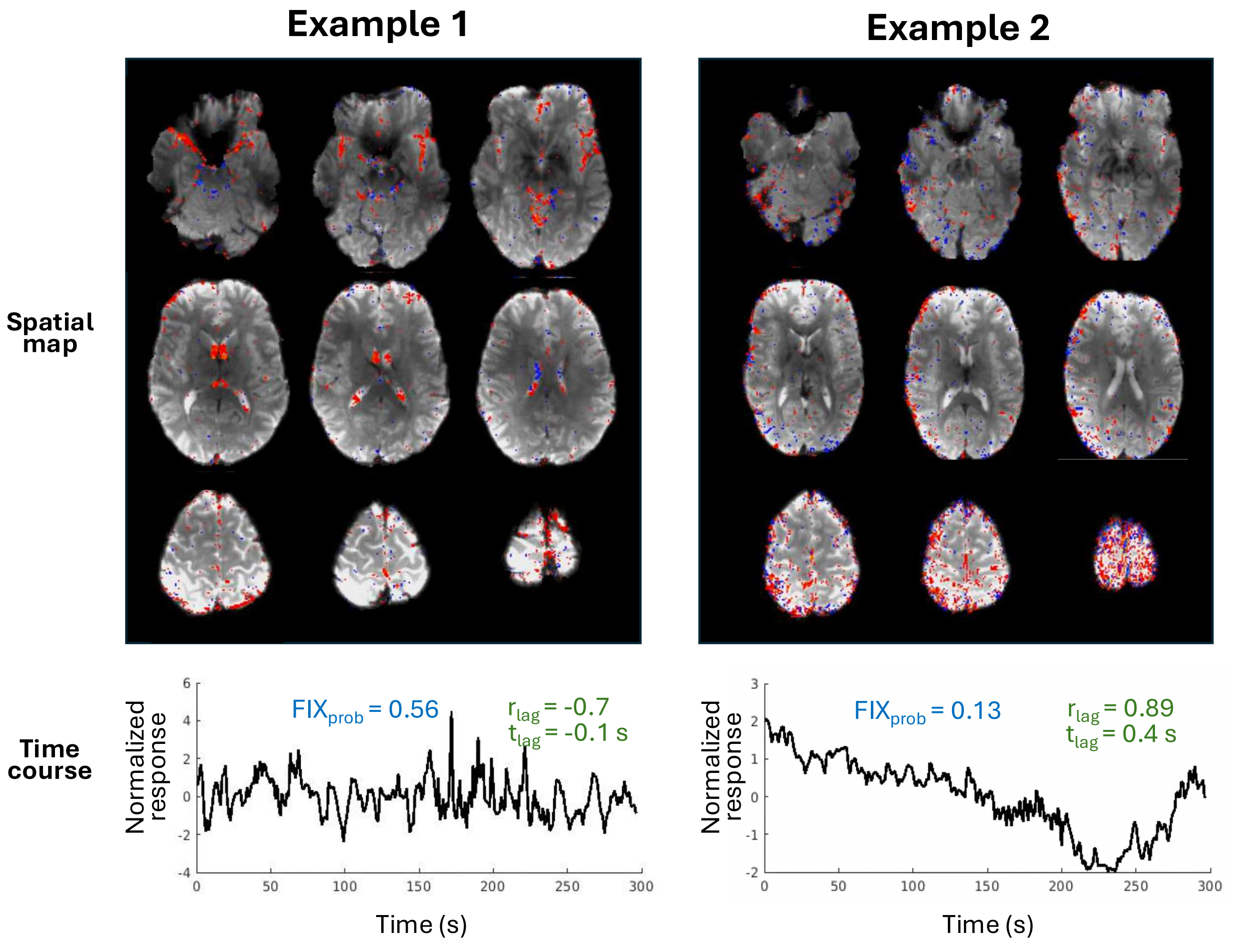


**Figure S6:** Example noise ICs misclassified as "signal" components either by FIX ('Example 1', FIX IC probability ($FIX_{prob}$) > 0.5) or by CortiLag-ICA ('Example 2', $r_{lag} \geq 0.2$ and $t_{lag} \geq 0.2$ s). The IC in 'Example 1' was visually identified as noise because the majority of its voxels were localized to the ventricles and large vessels; 'Example 2' was identified as noise because its spatial pattern (including voxels along the edge of the brain) and the time course (dominated by ultra-slow drifts) closely resembled motion artifacts. While there is general agreement between the classification provided by FIX-ICA and CortiLag-ICA, these examples demonstrate that neither classifier is perfect, and each misclassifies components that are clearly recognizable by eye as noise, but for different reasons.

### Supplementary Methods II: De-noising 3-mm isotropic visual task data using CortiLag-ICA

In addition to the moderate- and high-resolution fMRI data described in the main text, we additionally assessed the performance of CortiLag-ICA de-noising in a small, additional cohort of lower-resolution 3-mm isotropic fMRI data acquired at 7 Tesla[3] (five subjects, TR/TE = 745/26 ms, flip angle = 52°, FOV = 192×192 mm$^2$, 39 slices with no gap, acceleration factor $R$ = 3, multiband factor = 3, nominal echo spacing = 0.69 ms, bandwidth = 2368 Hz/pixel, with all other acquisition details identical to those of our main $N$=12 cohort). Each subject underwent a visual task scan (4 blocks, 16/24 s on/off per block), viewing black-and-white 'scaled noise' stimulus pattern flickering at 8 Hz during the 'on' condition, and neutral gray background during the 'off' condition[3]. The results are presented in Supplementary Figure S7 below.

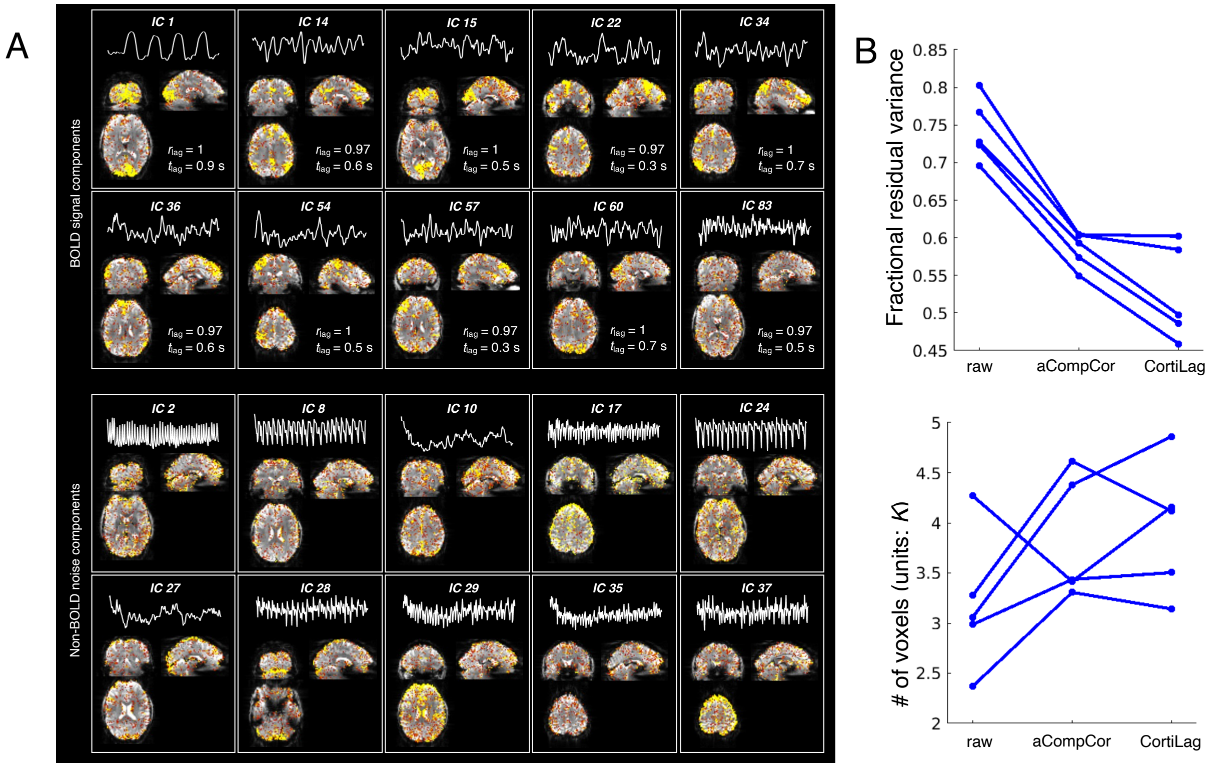


**Figure S7:** (A) In a representative subject, ICs with the highest $r_{lag}$ and $t_{lag}$ values (BOLD signal components) and low $r_{lag}$ and $t_{lag}$ values (non-BOLD noise components, all displayed components showed zero across-depth lags). Each panel shows the time course and the spatial pattern of each IC. Components with the highest $r_{lag}$ and $t_{lag}$ values comprised the task-active (IC 1) and common functional networks; whereas components with the zero cross-depth lags resembled typical artifacts. (B) Influence of de-noising on visual task activation: raw vs. aCompCor vs. CortiLag-ICA method. Both the mean squared error of residual variance (fraction relative to the total variance of each voxel's time series, correcting for degree of freedom changes) within the primary visual cortex, V1 (top; "Fractional residual variance") and task-active voxel counts (bottom; "# of voxels", $t$-score > 3) in the entire occipital cortex were quantified for comparison. "raw": fMRI data after basic preprocessing (rigid-body co-registration and slice-time correction); "aCompCor": including six motion parameters and five aCompCor regressors in the GLM analysis; "CortiLag": including noise ICs ($r_{lag}$ < 0.2 or $t_{lag}$ < 0.2 s) as nuisance regressors in the GLM analysis. Each blue line indicates the results of a single subject. These results suggest that CortiLag-ICA also holds promise in de-noising fMRI data at conventional spatial resolutions.
